# Perceived timing of postural instability onset

**DOI:** 10.1101/2023.03.16.533035

**Authors:** Robert E McIlroy, Michael Barnett-Cowan

## Abstract

**Background:** This study examines the perceived timing of postural instability onset, a crucial aspect of balance. Previous research using Temporal Order Judgment (TOJ) tasks found that postural perturbations need to occur significantly earlier than an auditory reference stimulus for individuals to perceive them as simultaneous. However, there are methodological concerns with this prior work, specifically regarding an unbalanced stimulus onset asynchrony (SOA) distribution.

**Research question:** Does the point of subjective simultaneity (PSS) between postural perturbation onset and an auditory reference stimulus differ for equal versus unequal SOA distributions?

**Methods:** A repeated measures design was utilized, presenting two different SOA distributions to 10 participants using a TOJ task during both the equal (72 trials) and unequal (88 trials) SOA distributions. Paired t-tests were used to determine if there was a significant difference between the PSS of the equal and unequal SOA distributions. Additionally, one-sample t-tests were performed on the PSS values of both the equal and unequal SOA distributions in comparison to 0ms (defined as true simultaneity) to determine if perceptual responses were delayed.

**Results:** The results indicate that the unequal SOA distribution resulted in a perceived delay of postural instability onset by 20.34ms, while the equal SOA distribution resulted in a perceived delay of the auditory stimulus of 3.52ms. However, neither condition was significantly different from each other nor from true simultaneity.

**Significance:** These findings suggest that the perception of postural instability onset is not slow, as previously thought, and highlight the importance of controlling methodological parameters when investigating sensory cues. This knowledge highlights the importance of controlling methodological parameters when investigating the perception of sensory cues and will help inform falls prevention strategies.

## 1. Introduction

Reliable perception of our external surroundings relies on the ability of the central nervous system (CNS) to rapidly detect self-motion and integrate information from different sensory modalities. This is not an easy task as the CNS must determine which information belongs together and which does not. Further, different modalities have varying transmission and processing speeds of each modality; this is known as the binding problem [1]. There has been significant research exploring the perceived timing of various sensory modalities, particularly auditory, visual, tactile, and vestibular [2-7]. This research has demonstrated that the perceived timing of sensory cues varies across modalities, and that differing temporal and spatial characteristics of modalities affects the perceived onset of stimuli. However, there is a paucity of research exploring the perceived timing of multisensory events that have a direct impact on safety and survival such as falls. Such research could provide valuable insights into the processing mechanisms of the CNS in fall-risk environments where fast and accurate sensory integration is critical. It is imperative that future research investigates the multisensory timing mechanisms in such contexts to enhance our understanding of the perceptual processing of multisensory events that have real-world consequences.

Our lab has recently conducted research investigating the perception of self-motion onset and postural instability using temporal order judgment tasks (TOJ). Our prior findings suggest that self-motion must precede a reference auditory stimulus for one to perceive them as occurring simultaneously, indicating that the perception of self-motion is slow [6-8]. Similar results were observed for the perceived onset of postural instability in both younger adults [9,10] and older adults [10]. Specifically, younger adults required falls to occur 44ms before a reference auditory stimulus to be perceived as occurring simultaneously [9,10], whereas this perceptual delay for falls was significantly longer in older adults (88ms; [10]). While slow perception of vestibular inputs [11] could explain this delay, here we assess an alternative hypothesis that the delay may stem from the methodology.

The TOJ task is a reliable psychophysical method for assessing the accuracy and precision of perceiving temporal events [12]. This task presents two stimuli with varying stimulus onset asynchronies (SOAs), and participants indicate which event occurred first. The point of subjective simultaneity (PSS; accuracy) and the just noticeable difference (JND; precision) can be determined from these responses. The PSS is the SOA at which participants are 50% likely to detect one stimulus before the other, while the JND is the interval between 50% and 75% [12]. Using a range of SOAs around true simultaneity (0ms), researchers can quantify an individual’s perception of the relative timing of two stimuli. Typically, SOAs are equally distributed around 0ms, and the method of constant stimuli is employed. In this method, the same number of trials are repeated at each SOA to obtain an average response at each SOA. This design reduces potential bias in participants’ responses and allows for reliable assessment of perceptual accuracy and precision.

Our prior work measuring the perceived timing of postural perturbations used an SOA distribution that was not precisely controlled. In these studies, postural perturbations were manually triggered in response to a visual “go” signal [9,10], resulting in a random distribution of SOAs. While this protocol has been used in other types of self-motion studies, the use of a manually triggered stimuli poses the risk of introducing unreliability and inconsistency in SOAs. In contrast, the method of constant stimuli is more commonly employed in TOJ tasks. With this method, fixed SOAs are repeated several times in a random order to obtain a more reliable measurement of an individual’s perception of the relative timing of two presented stimuli.

The present study aims to reassess the perceived timing of postural perturbations using a controlled and repeated SOA distribution. Previous studies have shown that the continuous presentation of asynchronous stimuli can directly influence the PSS during a TOJ task [13-17]. As cortical activity related to postural events occurs rapidly within the range of 90-150ms [18-21], perceptual delays for postural perturbation onset are questionable, especially as these fall-related cortical responses are comparable to those observed related to auditory events (∼50-150ms) [22-24]. Finally, reproducibility and falsifiability are essential aspects of scientific research [25,26], and as such, the present study seeks to replicate and extend prior work on the perceived timing of postural perturbations. The primary a priori hypotheses are that an equal SOA distribution will result in a mean PSS that is closer to true simultaneity compared to an unequal SOA distribution and that an equal SOA distribution will result in a mean PSS that is not significantly different from true simultaneity.

## 2. Methods

### 2.1. Participants

Ten healthy young adults (5 women; 21-26y) participated in this study. Participants reported no known musculoskeletal, auditory, visual, vestibular, or neurological disorders that may have affected their ability to perform the task. The study was conducted in accordance with the guidelines of the University of Waterloo Research Ethics Committee, and informed written consent was obtained from all participants after providing them with detailed information about the study procedures.

### 2.2. Protocol

The experimental setup and design is shown in Figure 1. This study used two different trial types: unequal SOA distribution and equal SOA distribution (Figure 2). Each of the outlined SOAs in Figure 1 was repeated eight times. Participants completed 160 trials (unequal = 88; equal = 72 trials). The timing of SOAs was confirmed by recording the auditory stimulus and load cell release.

**Figure 1:**
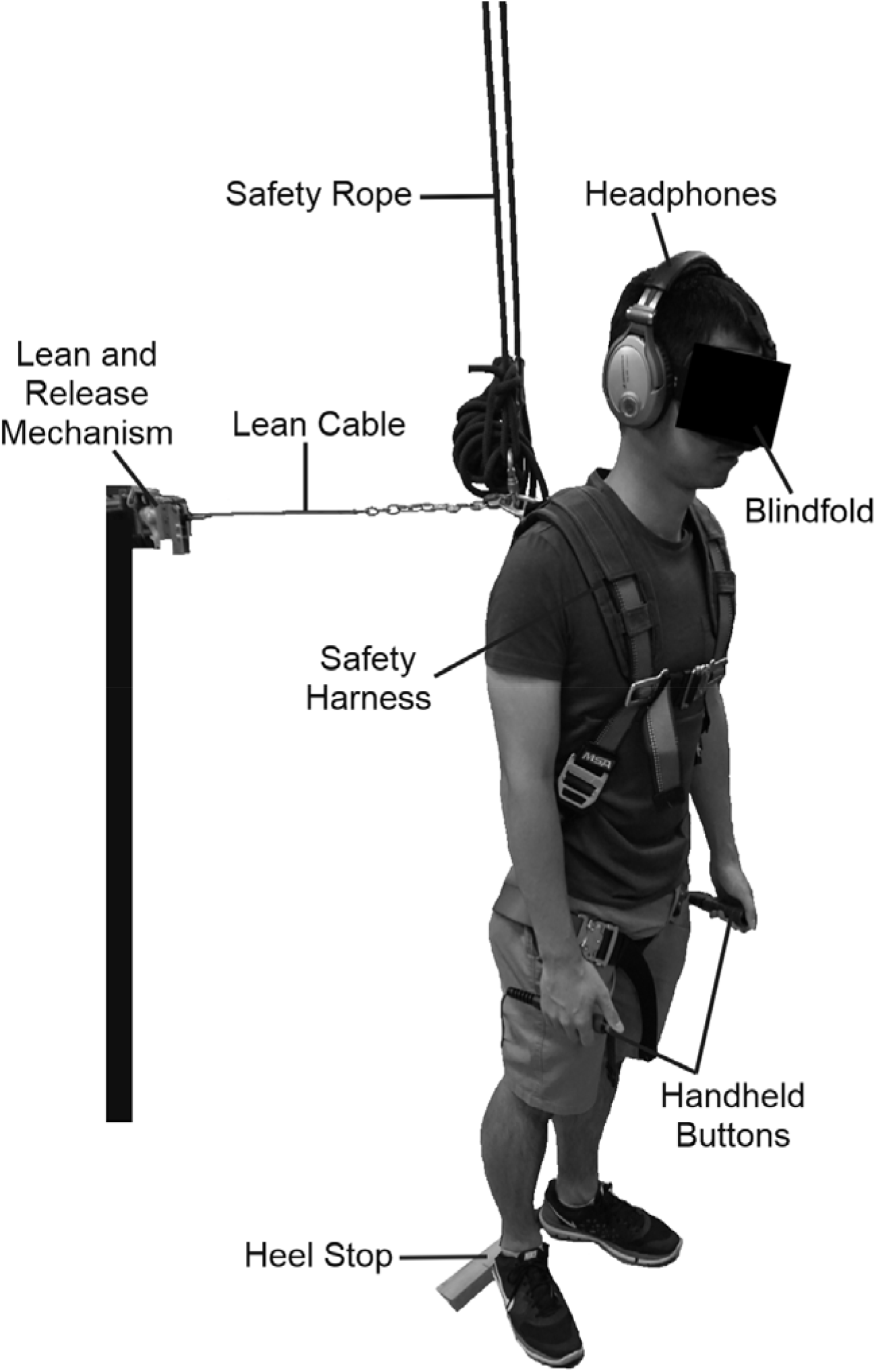
Participants were first weighed to determine the 7-8% lean angle to be utilized during the study. Participants were then fitted with a full-body harness that allowed for the attachment of a lean cable at the level of the 2nd and 3rd thoracic vertebrae and a safety rope, which was secured to the ceiling [9,10,27,28]. Participants were positioned ∼1m from the lean-and-release apparatus, and their feet were positioned in a standardized position [9,10,29]. Participants wore a blindfold through the duration of the experiment to remove visual cues. Postural perturbations were evoked by a lean-and-release mechanism and an auditory reference stimulus was produced through headphones. The postural perturbations produced a large enough perturbation to inherently evoke a stepping response in each of the participants [9,10,27,30,31]. Participants then made non-speeded TOJ responses with the use of handheld buttons (left button - auditory reference first; right button - postural perturbation first).

**Figure 2:**
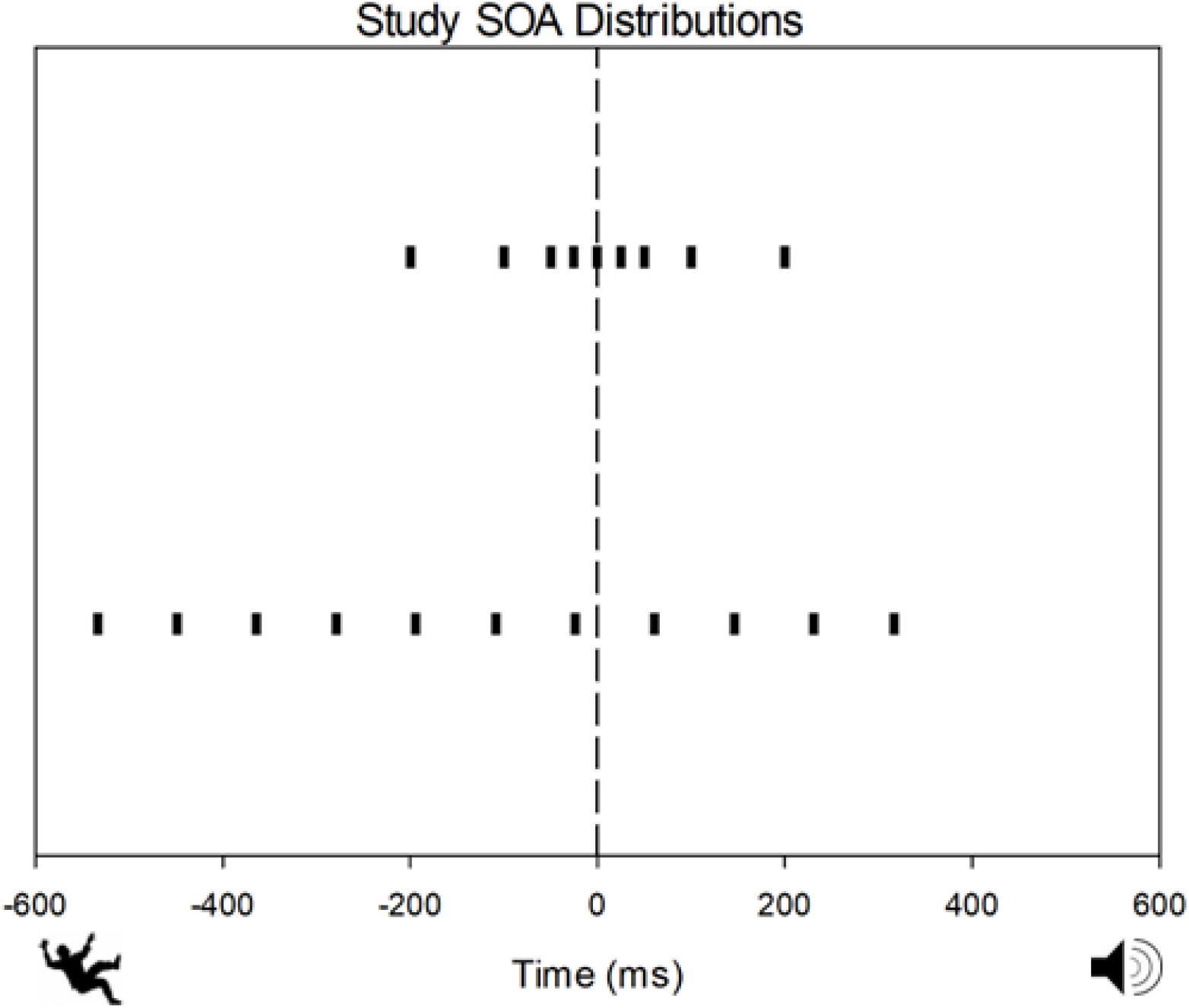
The equal (top) distribution consisted of the following SOAs: -200ms, -100ms, - 50ms, -25ms, 0ms, 25ms, 50ms, 100ms, 200ms. The unequal SOA distribution consisted of the following based on previous work in our lab [9]: -534ms, -449ms, -364ms, -279ms, - 194ms, -109ms, -24ms, 61ms, 146ms, 231ms, 316ms. Negative SOAs represent perturbation onset occurring first while positive SOAs represent the auditory reference occurring first. The equal SOA distribution at the top contains 4 SOAs in which the perturbation onset occurs first and 4 SOAs in which the auditory reference occurs first. The unequal SOA distribution at the bottom contains 7 SOAs in which the perturbation onset occurs first and 4 SOAs in which the auditory reference occurs first.

The unequal distribution of SOAs was utilized to replicate the SOA distribution used in previous research in the same laboratory [9]. To generate the unequal SOA distribution, minimum and maximum SOA values from each individual in the initial dataset were averaged and binned to 10% SOA increments, resulting in 85ms differences between SOAs. This distribution yielded a similar proportion of trials presenting postural instability first as in previous research [9]. Unlike prior work, each SOA was repeated an equal number of times in this study. The equal distribution of SOAs surrounding true simultaneity was set between - 200ms-200ms, as SOAs exceeding 200ms are distinguishable by participants according to prior research [2,9,10]. The SOAs were subsequently reduced by 50% until a central value of 0ms was reached, passing through values of 25ms and -25ms. The presentation of both unequal and equal SOA distributions was counterbalanced across participants, and SOAs were fully randomized for each participant. For each 15-20s trial, participants kept their arms and hands comfortably at their sides while leaning at the appropriate angle. Participants completed five practice trials.

### 2.3. Lean-and-Release Apparatus & Stimuli

Figure 3 depicts the crossbow release mechanism that was mounted on the load cell and held a lean cable that was fastened to the participant via a harness. In contrast to our prior studies [9,10] where a manual release method was employed, an automated technique was adopted in the present study to allow for precise perturbation onset times.

**Figure 3:**
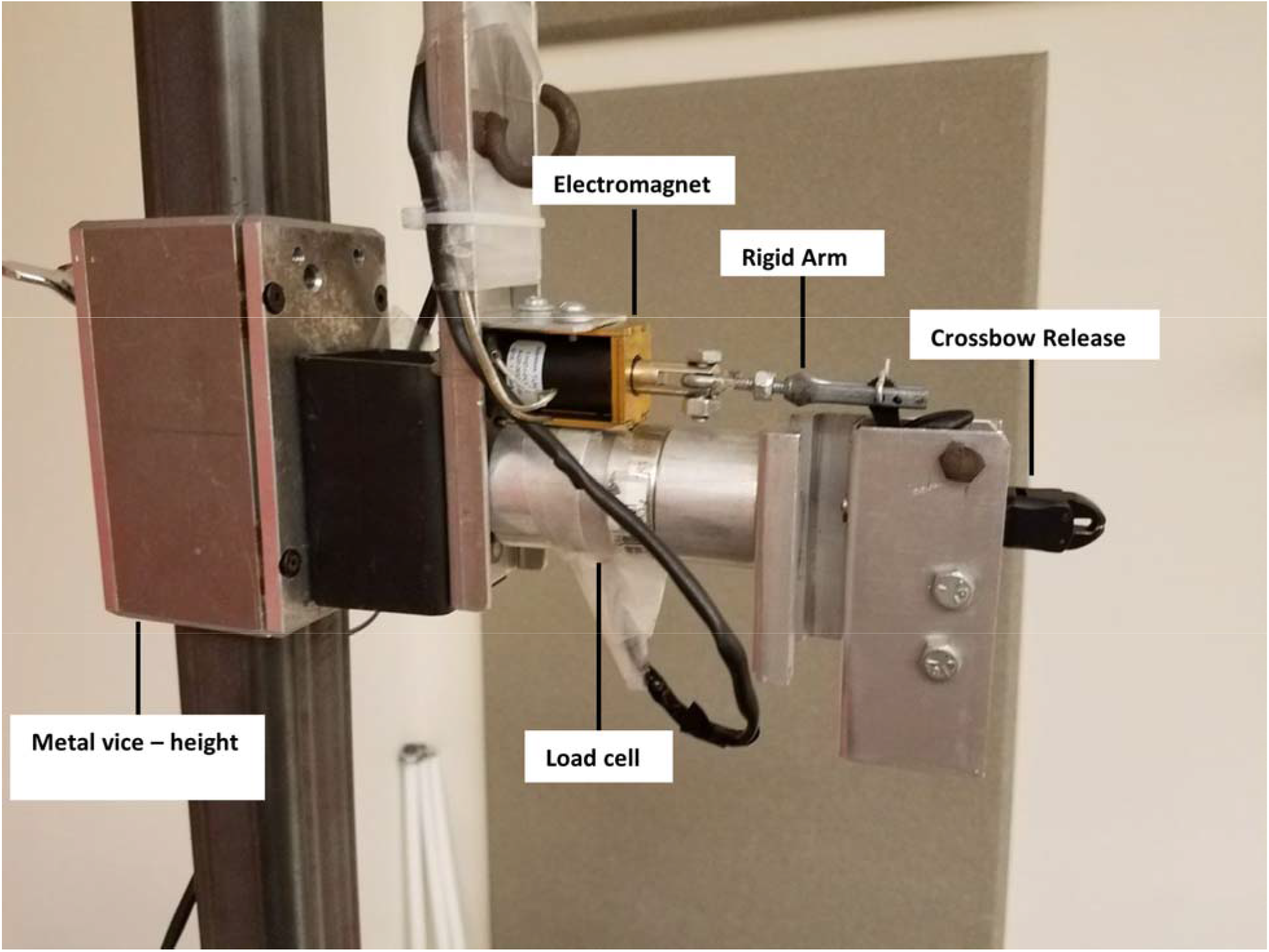
The lean-and-release apparatus consisted of three components: a crossbow release, a load cell, and an electromagnet. The crossbow release, load cell and electromagnet were attached to a metal vice which could be vertically moved to adjust for each participant’s height. The load cell was capable of measuring 454kg of force and sampled at 100 Hz. Load cell readings determined the onset of postural perturbation and ensured that participants maintained a lean angle between 7-8% of their body mass, which was close to the target of 10% [27,30,31]. However, the crossbow release was unable to consistently release for individuals who exceeded certain weights. The electromagnet was utilized to activate the crossbow release. The electromagnet was connected to the crossbow release arm via a rigid metal piece to minimize initiation variability. Activation of the electromagnet was triggered by a 5V pulse sent from a LabVIEW (National Instruments, Austin, Texas, USA) program, causing the rigid metal arm to open the crossbow release and initiate the postural perturbation.

### 2.4. Auditory Stimuli

The LabVIEW program that produced the 5V pulse to the electromagnet also generated the auditory reference stimulus. A 250ms 500Hz square wave soundburst was delivered via noise-canceling headphones (Sennheiser PXE 450) [c.f. 9,10]. The volume of the white noise, which was constantly present throughout the study via a free cell phone application (White Noise Free), was manually adjusted such that each participant could not hear external noise from the lean-and-release mechanism. This ensured that participants were not receiving additional auditory cues that could alert them about the impending postural perturbation, while still allowing them to distinctly detect the onset of the auditory reference. The 500Hz auditory reference stimulus was well above the threshold, ensuring that individuals could clearly detect the auditory reference above the white noise.

### 2.5. Data Analysis

Binary TOJ responses were assigned 0 for “postural perturbation first” and 1 for “auditory reference stimulus first”. Average TOJ responses for each SOA were plotted as a probability of the sound occurring first. Negative SOAs represent that the postural perturbation occurred first and positive SOAs represent that the auditory reference occurred first. A two parameter logistic function (Eq. 1) was fitted to each of the participants’ averaged responses as a function of SOA using SigmaPlot 12.5.

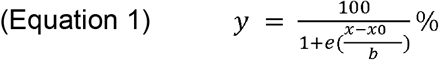

Where the inflection point of the logistic function (x_0_) was taken as the point of subjective simultaneity (PSS) and the standard deviation (b) was taken as the just noticeable difference (JND) [9].

Outcome parameters were inspected to identify potential outliers or poor quality fits. If both the R^2^ of the logistic regression fit was ≤ 0.5 [32] and the *p*-value was *p* ≥ 0.05 then we would remove the participant’s data from the analysis as we could not be confident that the logistic regression provided an accurate fit.

### 2.6. Statistical Analysis

We used a within-subjects’ design to assess the hypothesis that postural instability will not be perceived significantly slower than the auditory reference. A one-sample t-test was conducted comparing the mean PSS value of the condition to true simultaneity (0ms). The one-sample t-tests specifically assessed whether the mean PSS values were significantly different from zero. To investigate the effect of the distribution on mean PSS, a paired sample t-test was performed to compare the Unequal and Equal distribution conditions. If normality failed in any comparisons as assessed by the Shapiro-Wilk test, then there was an attempt to normalize the data. If these attempts with Z-Score normalization were not successful, then non-parametric statistics were performed. For each of the statistical tests, a significance level of □ = 0.05 was utilized.

## 3. Results

No individuals violated the conditions that were set for this study. All participants’ data when fitted with the non-linear regression logistic function exhibited R^2^ values ≥ 0.5 and a *p*-value ≤ 0.05. PSS and JND values of each individual were derived from logistic regressions that were fit to the averaged responses at each SOA. These fits for the unequal and equal distributions are depicted in Figure 4. Similar to our lab’s prior work [9,10], the PSS of the unequal distribution was negative (mean = -20.34ms, SE = 18.67, median = -1.17ms; Figure 5), indicating that on average the onset of the postural instability needed to occur prior to the auditory reference to be perceived as simultaneous. The equal distribution produced a positive PSS (mean = 3.52ms, SE = 11.97, median = 3.27ms; Figure 4), indicating on average the auditory reference needed to occur prior to the postural instability to be perceived as simultaneous. However, neither the unequal (One Sample Signed Rank Test (one tail); Z = - 0.459, *p* = 0.348) nor the equal (One Sample T-test (one tail); t(9) = 0.294, *p* = 0.388) PSS values were significantly less different than 0ms (true simultaneity). This supports our initial hypothesis that individuals would not perceive the onset of postural instability slower than the auditory reference stimulus. When directly comparing the unequal and equal distribution PSS values, there was an observed difference of 23.86ms, however this difference was not statistically significant (Wilcoxon signed rank test; Z = -1.274, *p* = 0.232, 1 - □ = 0.24).

**Figure 4:**
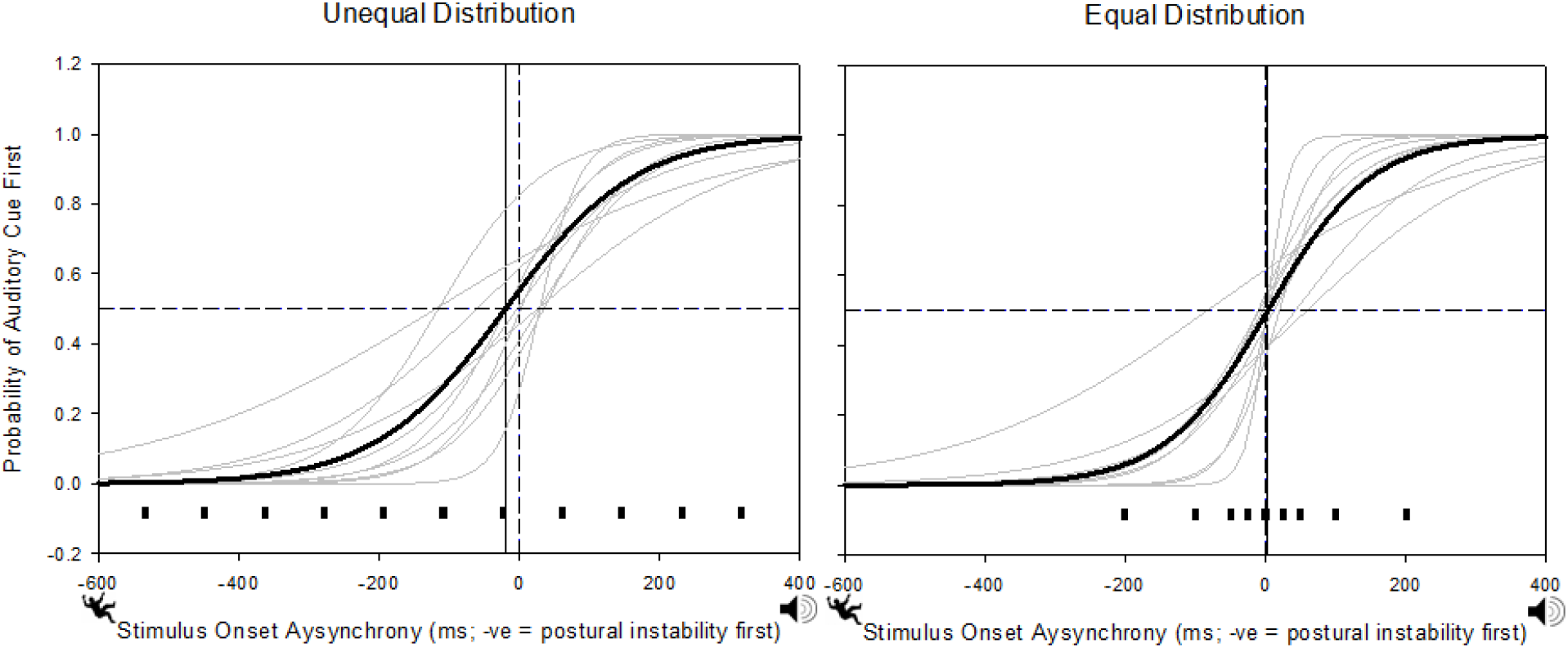
Logistic fits for individuals (light grey) and averaged (black) responses in each TOJ task. The solid vertical lines represent the mean PSS value for each condition. The average PSS from the unequal distribution was -20.34ms and the average PSS from the equal distribution was 3.52ms. An SOA of 0ms represents true simultaneity and is indicated by the vertical dashed line. The horizontal dashed line represents 50% probability of the auditory stimulus perceived as occurring first. The small black hash marks at the bottom of the graphs represent the SOAs for the unequal distribution (left) and equal distribution (right).

**Figure 5:**
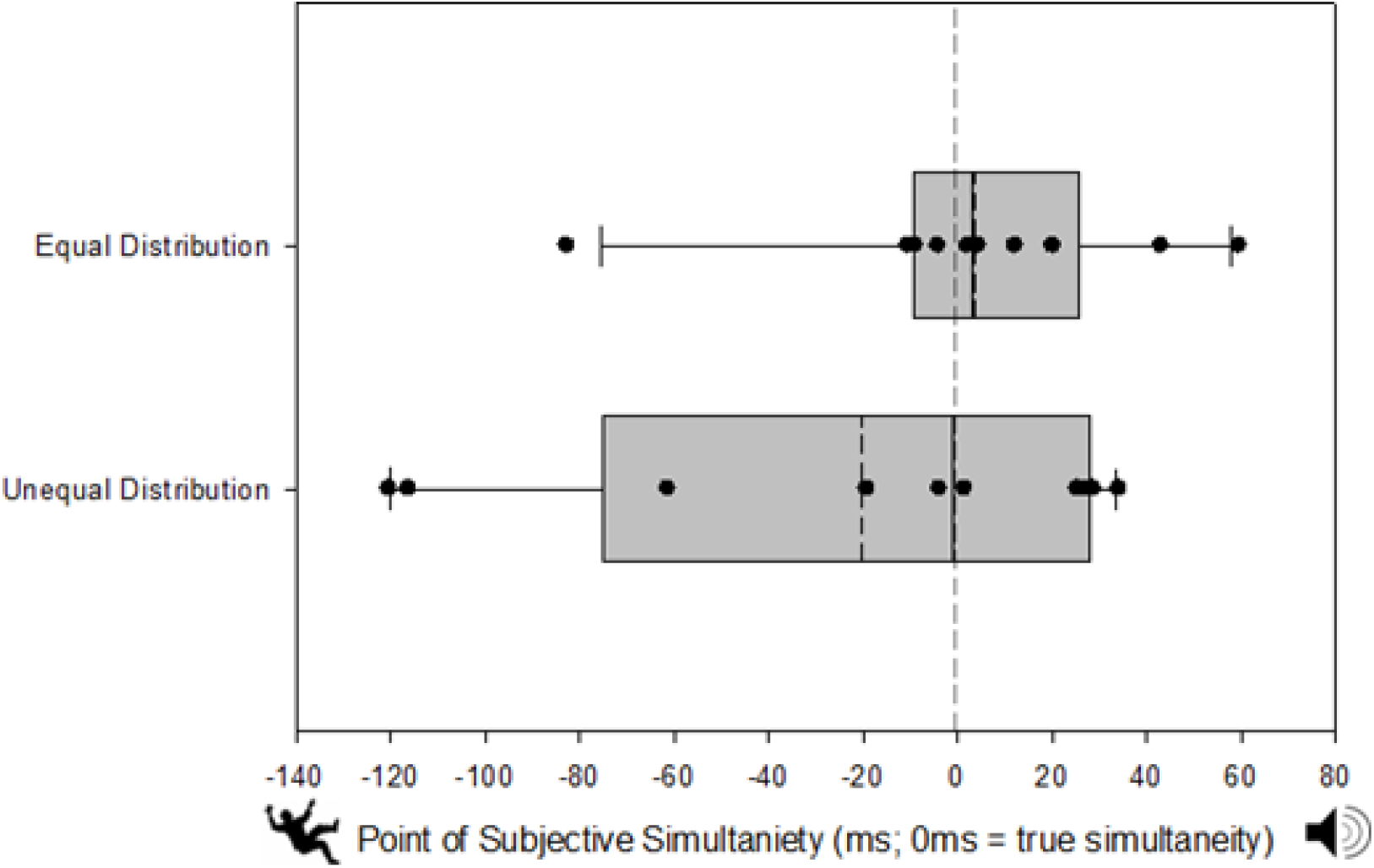
Mean PSS (dashed line), median PSS (solid line), and individual PSS values (circles) for the equal (top) and unequal (bottom) SOA distributions. Grey bars represent the 25-75th percentiles of the PSS data with error bars representing the 90th and 10th percentiles of data distributions. There were no statistically significant differences between the unequal and equal distribution PSS values.

The current study did not have an a priori hypothesis regarding JND values. The JND values for the unequal (mean = 92.86ms, SE = 16.42, median = 75.76ms) and equal (mean = 73.82ms, SE = 15.51, median = 66.08ms) distributions had a mean difference of 19.04ms, but were not statistically significant from each other (Paired Sample T-test; t(9) = -1.373, *p* = 0.203, Figure 6).

**Figure 6:**
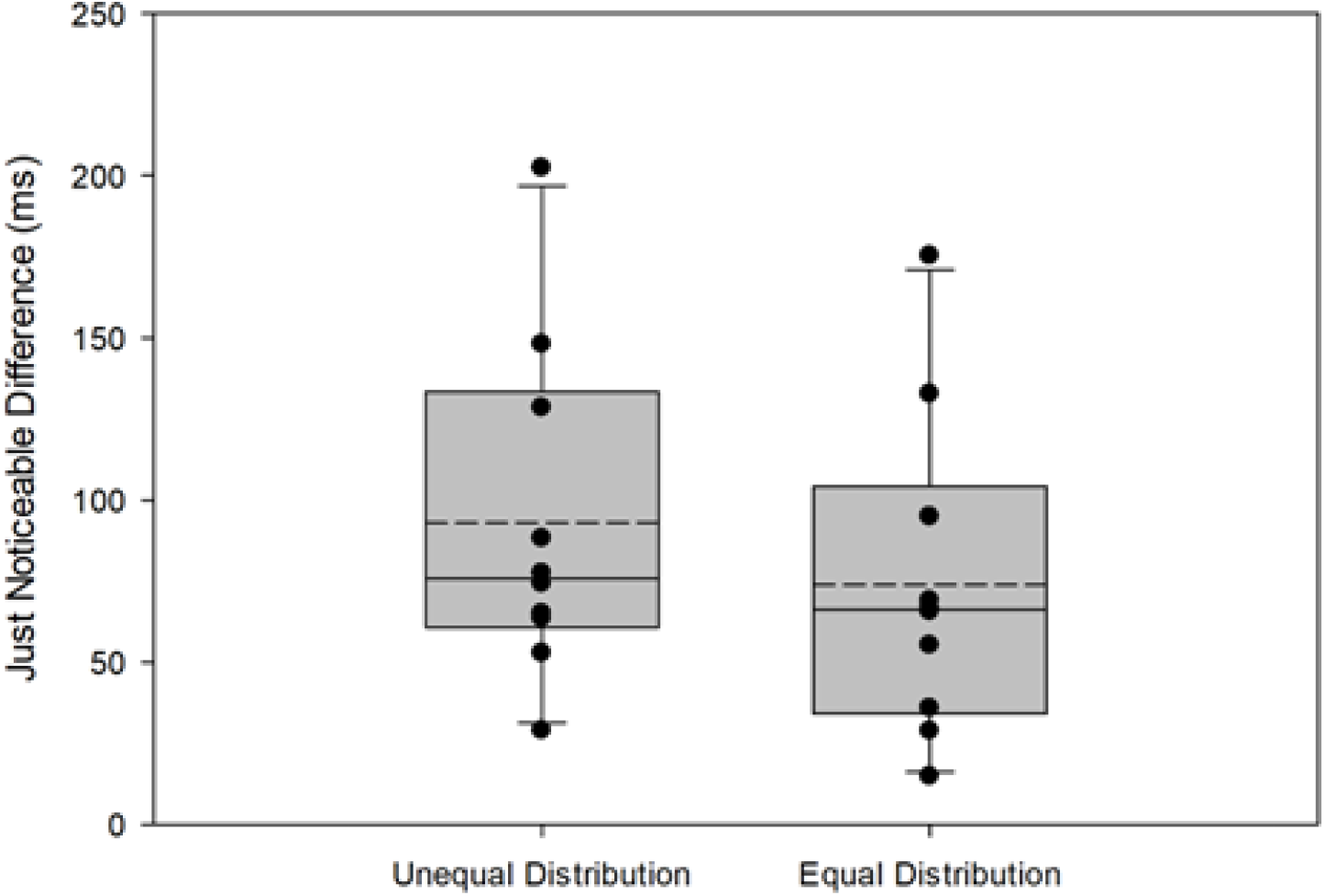
Mean PSS (dashed line), median PSS (solid line), and individual JND values (circles) for the equal (top) and unequal (bottom) SOA distributions. Grey bars represent the 25-75th percentiles of the JND data with error bars representing the 90th and 10th percentiles of data distributions. There were no statistical differences between the unequal and equal distribution JND values.

Comparing PSS results from two of our prior works [9,10] (mean = -44.16ms, SE = 8.23, median = -44.00ms, respectively) to the present unequal distribution PSS results, the difference of 23.82ms was not statistically significant (Mann-Whitney Rank Sum Test; U = 62.00, *p* = 0.099). However, compared to the present equal distribution PSS results, the difference of 47.27ms was statistically significant (Independent Sample T-test; t(28) = 3.315, *p* = 0.0025). Thus, we were able to confirm that only when a manual random distribution of SOAs is used, the postural perturbation needs to precede the sound reference stimulus to be perceived as simultaneous [c.f. 9,10].

## 4. Discussion

Reproducibility and falsifiability are essential aspects of scientific research [25,26]. Prior work in our lab found that a postural perturbation needs to occur approximately 45ms prior to an auditory reference stimulus in order to be perceived as simultaneous, suggesting that the perceived timing of falls may be slow [9,10]. This study investigated how TOJ task design characteristics impact equal and unequal SOA distributions effects on the PSS. Our results demonstrate that the PSSs for unequal and equal distributions were not significantly different from each other or from true simultaneity (0 ms). These new found results suggest that the perceived timing of falls is not slow compared to an auditory reference stimulus.

Equal SOA distributions and the method of constant stimuli are commonly used in TOJ tasks [2,5,14,32-34] to avoid biasing the PSS [20-24]. When SOA distributions are biased in one direction, prior work has shown that the PSS shifts in this same direction 13-17]. Miyazaki et al. examined the effects of a shifted Gaussian distribution on the PSS during tactile and audiovisual TOJ tasks [13]. They found that the PSS shifted away from the peak Gaussian distribution during tactile-tactile TOJ, possibly due to Bayesian calibration/integration, and towards the peak Gaussian distribution during audiovisual TOJ, possibly due to lag adaptation/temporal adjustment. Bayesian calibration/integration refers to the use of prior information to improve or impact sensory estimates, while lag adaptation/temporal adjustment describes the shift in perception due to consistent exposure to asynchronous stimuli.

In this present study, the average unequal distribution SOA was -106ms (favouring postural perturbation prior to the audio reference stimulus), while the equal distribution was centered on 0ms. In our lab’s prior work [9,10], the average SOA could not be defined due to the random presentation of stimuli. Nevertheless, assuming the unequal SOA distribution used in our study as an analog to this prior research, we estimate that the SOA was also around -106ms. With an SOA shift of -106ms, the lab’s prior studies [9,10] produced a point of subjective simultaneity (PSS) of -44ms, whereas our results show that the PSS was -20.34ms, and with an SOA centered on 0ms, the PSS was 3.52ms. Our findings, together with the lab’s prior studies [9,10], suggest that the unequal SOA distribution evoked lag adaptation or temporal adjustment. However, it is important to note that in the present study, the median value for the unequal distribution was closer to 0ms (−1.17ms), indicating a weaker effect of lag adaptation. Miyazaki et al [13] proposed that lag adaptation or temporal adjustment is present in audiovisual tasks due to differences in the detection, transduction, and processing of auditory and visual information. Our study utilized an auditory reference and a multisensory event (postural instability) that affected the proprioceptive/somatosensory and vestibular systems, rather than the visual system, since participants had their eyes closed. Therefore, it is feasible that lag adaptation or temporal adjustment was affecting the perceptual responses in the unequal SOA distribution in comparison to the equal SOA distribution.

Sample size is an important factor to consider as a limitation of the present and past studies [9,10]. The power of the comparison between equal and unequal SOA conditions was 1 - □ = 0.24, indicating a higher probability of type II error in our statistical analysis. With an a priori power analysis, a population of 67 participants would be required to achieve a power of 1 - □ = 0.95. Thus while the sample size that we used for the present study was chosen to replicate a previous study that only recruited eight participants [9]. A larger sample size could have led to greater normalization of variability observed in the population. Therefore, the sample size must be considered as a potential limitation of this study. Individual differences must also be considered as they play a significant role in perceptual outcome measures of postural instability onset and auditory reference onset (see Figures 4 & 5). Variability in individual responses has been observed in perceptual tasks [16,34,35]. Attention could also be a potential factor for individual differences, as attention towards one stimulus over the other has been observed to shift the perception of simultaneity towards the attended stimuli [5,16,36]. Although participants were encouraged to attend to the perturbation onset and sound onset equally, it is still possible that some individuals attended more to one stimulus than the other.

The findings of this study suggest that a more controlled methodology may eliminate the previously observed perceptual delay in postural instability onset. These results highlight the importance of carefully designing and implementing research methods, as they can directly impact the outcome measures. Specifically, the results demonstrate how the use of different SOA distributions can affect perceptual outcome measures, such as the PSS. A controlled methodological design enables greater consistency across participants, and enhances the reproducibility of the findings. Consistency across participants is critical to obtaining meaningful results and making valid comparisons with other research. Moreover, researchers should carefully consider potential methodological differences when comparing their results to those of other studies, as variations in equipment, stimuli, and parameters could significantly affect the results.

## Conflict of Interest

There are no conflicts of interest.

## Acknowledgements

Supported by a Natural Sciences and Engineering Research Council of Canada (NSERC) Discovery Grant and an Ontario Early Researcher Award to MB-C. Thanks to Jeff Rice for helping assemble the lean-and-release mechanism.

## Notes

### Competing Interest Statement

The authors have declared no competing interest.

